# The bacterial microbiota of a parasitic plant and its host

**DOI:** 10.1101/775155

**Authors:** Connor R. Fitzpatrick, Adam C. Schneider

**Affiliations:** Department of Biology, University of Toronto Mississauga, Mississauga, ON, Canada L5L 1C6; Department of Integrative Biology, University of California, Berkeley, California, USA, 94720; Department of Biology, University of North Carolina, Chapel Hill, North Carolina, USA, 27599; Department of Biology, Hendrix College, Conway, Arkansas, USA, 72032

**Keywords:** holoparasite, network analysis, Orobanchaceae, parasitic plant, plant microbiome, Procrustes, microbiome assembly, *Orobanche hederae*

## Abstract

How plant-associated microbiota are shaped by, and potentially contribute to the unique ecology and heterotrophic life history of parasitic plants is relatively unknown. Here, we investigate the leaf and root bacterial communities associated with the root holoparasite *Orobanche hederae* and its host plant *Hedera* spp. We sequenced the V4 region of the 16S rRNA gene from DNA extracted from leaf and root samples of naturally growing populations of *Orobanche* and infected and uninfected *Hedera*. Root bacteria inhabiting *Orobanche* were less diverse, had fewer co-associations, and displayed increased compositional similarity to leaf bacteria relative to *Hedera*. Overall, *Orobanche* bacteria exhibited significant congruency with *Hedera* root bacteria across sites, but not the surrounding soil. Infection had localized and systemic effects on *Hedera* bacteria, which included effects on the abundance of individual taxa and root network properties. Collectively, our results indicate that the parasitic plant microbiome is derived but distinct from host plant microbiota, exhibits increased homogenization between shoot and root tissues, and displays far fewer co-associations among individual bacterial members. Host plant infection is accompanied by modest changes of associated microbiota at both local and systemic scales compared with uninfected individuals. Our results provide insight into the assembly and function of plant microbiota.

## Introduction

Plants harbour rich assemblages of microorganisms, which vary in diversity and composition across host plant tissues, individuals, and species (Bulgarelli *et al.*, 2013). This variation is driven by innate plant immunity (Lebeis *et al.*, 2015; Stringlis *et al.*, 2018), the quality and quantity of plant-derived resources (Zhalnina *et al.*, 2018), and microbe-microbe interactions (Agler *et al.*, 2016; Durán *et al.*, 2018). Controlled experiments indicate that microbiota may play a role in plant nutrient acquisition (Castrillo *et al.*, 2017) and tolerance to abiotic and biotic stress (Fitzpatrick *et al.*, 2018), including defense from pathogens (Vogel *et al.*, 2016). However, host plant species vary in the composition of their associated microbiota and the causes and ecological consequences of this variation are poorly understood (Fitzpatrick *et al.*, 2018). Increased insight into the assembly and function of plant microbiota requires investigation of plant species that occupy diverse ecological niches (e.g. Angel *et al.*, 2016; Coleman-Derr *et al.*, 2016; Finkel *et al.*, 2016). Such investigation provides an opportunity to address how associated microbiota are shaped by, and potentially contribute to the functional diversity found across plant species.

One of the most dramatic niche shifts undergone by plants is the transition from autotrophy to heterotrophy. In heterotrophic plants, resource acquisition from photosynthetic plants occurs indirectly through co-associated mycorrhizal fungi (mycoheterotrophy), or directly through specialized parasitic organs called haustoria. The haustorium attaches to the root or stem vasculature of host plants and acts as a conduit through which material is exchanged primarily from host to parasite (Sareendenchai and Zidorn 2008; LeBlanc *et al.*, 2012; Spallek *et al.*, 2017). Approximately 2% of flowering plants directly parasitize other plants to some degree in order to acquire fixed carbon, water, and/or mineral resources. Complete loss of photosynthetic function (i.e. holoparasitism) has evolved at least 13 times in flowering plants alone (Westwood *et al.*, 2010; Braukmann *et al.*, 2013; McNeal *et al.*, 2013). Unsurprisingly, this life-history shift has been accompanied by dramatic ecological and physiological shifts, as well as morphological and genomic changes collectively referred to as ‘parasitic reduction syndrome’ (Colwell *et al*., 1994). For example, root holoparasites typically exhibit vestigial, scale-like leaves, loss of a developed root system (Fig. 1A-C), and reduction in chloroplast genome size. However, very few studies have investigated changes induced in the composition and assembly of parasitic plant-associated microbial communities (Kruh *et al*., 2017) in spite of the mounting evidence that such communities are crucial for plant function and fitness (Vogel *et al.*, 2016; Castrillo *et al.*, 2017; Fitzpatrick *et al.*, 2018).

**Figure 1.**
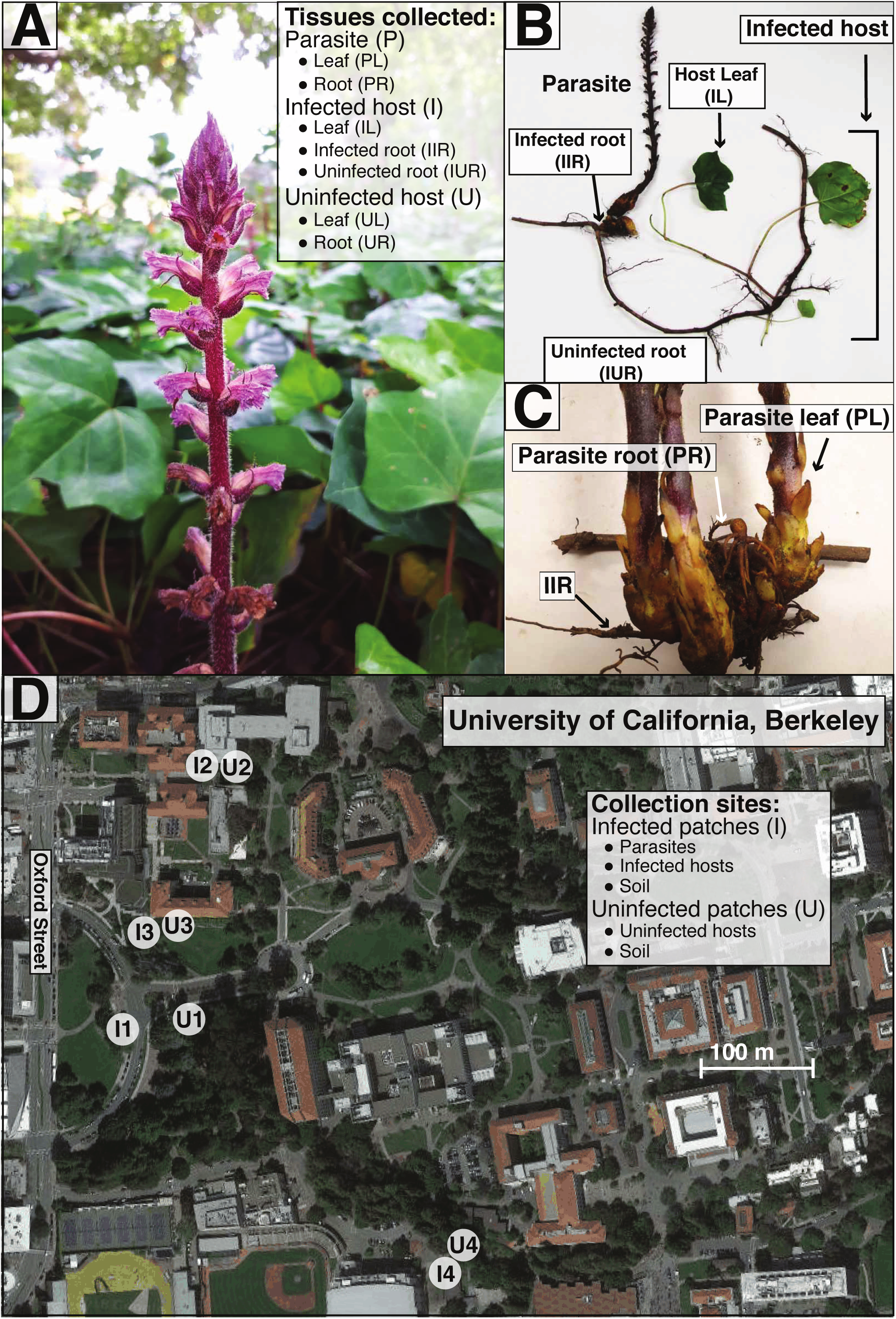
An overview of the study system and sampling design. (A) We sampled leaves and roots from *Orobanche* (inflorescence in foreground) and from infected and uninfected *Hedera* (green plant in background). Note that the abbreviations used here are used throughout the text. (B) We carefully excavated individual *Orobanche* and its *Hedera* host to sample leaves and roots from both organisms. For infected *Hedera*, we distinguished between infected roots (IIR), which were physically attached to the parasite, and uninfected roots (IUR), which did not exhibit direct physical attachment to any parasite. For uninfected *Hedera*, we sampled leaves and roots in a similar fashion. (C) We sampled sampled leaves (PL) and roots (PR) for *Orobanche*, which are homolgous but lack the functionality of leaves and roots found in non-parasitic plants. (D) We sampled plants and soil from paired infected and uninfected ivy patches at four sites on the University of California, Berkeley campus. From infected patches we collected *Orobanche* and *Hedera* samples and from uninfected patches we collected only *Hedera* samples.

Parasitic plants may also shape the microbiota of their host. Infected host plants have greater demands on their resources because of the significant portion of fixed carbon, water, and nutrients siphoned by the parasitic plant (Fernández-Aparicio *et al.*, 2016). Consequently, infected host plants mount physiological and molecular responses to resist the invading parasite. This can occur over both local scales (i.e. directly parasitized tissue) and systemic, whole plant scales (Hiraoka *et al.*, 2009). These defense responses include the production of reactive oxygen species (Hegenauer *et al.*, 2016) and defensive secondary compounds (Serghini *et al.*, 2001; Castillejo *et al.*, 2009), as well as the activation of hormonal pathways governed by jasmonic acid and ethylene (Hiraoka *et al.*, 2009). Plant immunological responses can re-structure associated microbiota due to variation in the susceptibility of microbial taxa to plant immune outputs (Lebeis *et al.*, 2015; Stringlis *et al.*, 2018; Voges *et al*, 2019). Finally, parasitic plant-derived molecules can alter host root growth (Spallek *et al.*, 2017), which could alter the composition of associated microbiota. Thus, by altering the quality and quantity of plant-derived resources available for microbes, activating innate plant immunity, and altering host plant morphology, parasitic plant infection could perturb the resident microbiota of their hosts.

In this study we address three main research questions. First, what are the differences in the diversity, composition, and structure of plant microbiota between a root holoparasite and its autotrophic plant host? We predicted that parasitic plants would exhibit reduced microbial diversity and altered composition and structure relative to host plants potentially due to the ‘parasitic reduction syndrome’. Second, does the composition of parasitic plant microbiota track the composition of host plant microbiota? Due to the biased flow of material through the haustorium from host to parasite we predicted that parasitic plant microbial composition would exhibit congruency with that of its host. Third, does infection by a parasitic plant result in perturbations to host plant microbiota? We predicted that parasitic plant infection would alter the diversity and composition of host plant microbiota, possibly a result of induced physiological and immune host responses, which are known to enrich for particular microbial taxa (Lebeis *et al.*, 2015; Stringlis *et al.*, 2018; Voges *et al*, 2019). To address these questions, we investigated the plant-associated bacterial microbiota from stable natural populations of *Orobanche hederae,* the ivy broomrape (hereafter *Orobanche*), and infected and uninfected individuals of its host plant, *Hedera* spp., ivy (hereafter *Hedera*).

## Materials and Methods

### Study System

*Orobanche hederae* and various species of *Hedera* are naturalized on the University of California Berkeley (UCB) campus, but native to Eurasia and North Africa. On the UCB campus, *O. hederae* was first collected in 2000 and has been persistent at numerous sites for well over 15 years (vouchers at UC and Jepson Herbaria), growing exclusively in near-monocultures of *Hedera algeriensis* or, rarely, *H. helix*. Curiously, the UCB campus is the only record of *O. hederae* outside its native range with the possible exception of a depauperate specimen collected in 1950 in Raleigh, NC, but not observed subsequently (Musselman, 1980). Similar to other root holoparasites, *O. hederae* seeds germinate in the presence of a suitable host plant and the seedling immediately attaches to a single host root, where it persists for the remainder of its life (Joel *et al.*, 2013). Organs homologous to the leaves and roots of photosynthetic plants are greatly modified in *O. hederae*: scale-like leaves and short, stout hairless roots composed of large, starch-containing cells surrounding the central vascular bundle (Tate, 1925; Joel *et al.*, 2013).

### Plant and soil sampling

We sampled from 4 sites across the UC Berkeley campus (Fig. 1D). At each site we located one infected and one paired but noncontiguous uninfected patch of ivy, assessed by a thorough visual inspection. Infected ivy patches possessed between 20 – 50 live *Orobanche* stalks as well as several dozen senesced stalks from previous years, whereas in uninfected patches we were unable to find any living or senesced parasite individuals. Thus, uninfected individuals in our study may represent resistant *Hedera* genotypes but may also represent individuals that have not been confronted with *Orobanche*. Infected and uninfected patches are consistent from year to year (A.C.S., pers. obs.). At each infected patch we carefully excavated flowering *Orobanche* individuals and the connected host plant root, and followed it until we located a connected leaf (Fig. 1B). The parasite and associated host, including aboveground and belowground organs comprised a single specimen (Fig. 1B). We repeated this process 3 times in each infected patch, sampling individuals at least 2 m apart. For each uninfected patch we collected three ivy individuals at least 2 m apart, which included aboveground and belowground tissues. Finally, we collected two soil samples (approximately 1 g sampled from 1-10 cm soil depth), from each infected and uninfected patch at each site.

After collection at each site (approximately 2 hours) plant samples were immediately brought back to the lab in separate bags and prepared for DNA extraction of endophytic microbes using an established protocol (Supporting Information: Methods). From infected patches, we harvested samples from parasite leaves and roots (PL and PR, respectively; Fig. 1C), fully expanded ivy leaves (IL), and two ivy root samples (Fig. 1B): infected roots (IIR; i.e. the specific ivy root being parasitized); and uninfected roots (IUR; i.e. roots from the same ivy individual and within 50 cm of the parasite, but not directly being parasitized). From uninfected patches, we harvested ivy leaves and roots (UL and UR, respectively). All samples were standardized by fresh weight and carefully selected to represent homologous organs between the parasite and host plant (full details in Supplementary Data: Methods). Voucher and host information is provided in Table S1.

### DNA Extraction and Sequencing

We used DNeasy PowerSoil extraction kits (Qiagen) following the manufacturer’s protocol to isolate DNA from plant tissue and soil samples, then amplified in triplicate, and sequenced the V4 region of the 16S rRNA gene using Illumina MiSeq 2 x 250 bp. Individual fastq files are archived on the NCBI Sequence Read Archive (SRP154488). For a detailed description of PCR and library preparation see Supplementary Data: Methods.

### Bioinformatics

We processed sequences using the R package ‘DADA2’ v. 1.8.0 (Callahan *et al.*, 2016) (Supplementary Data: Methods). DADA2 infers unique bacterial taxa from amplicon sequence variants (ASVs). We assigned taxonomy to individual ASVs and built a maximum likelihood phylogenetic tree from aligned ASV sequences. Using the R package ‘phyloseq’ v. 1.24.0 (McMurdie and Holmes, 2012), we removed ASVs that were unassigned to a phylum of Eubacteria (268 ASVs removed), or assigned to plastid and mitochondrial lineages (3268 ASVs removed), which left 6.5 million reads distributed across 18,329 ASVs. Our final dataset consisted of 103 unique microbiome samples (60 *Hedera* and 24 *Orobanche* samples [12 for each of the sample types shown in Fig. 1A], 15 soil samples, and 3 controls), each with on average 63,527 high quality sequences (± 5365; standard error).

### Statistical analyses

#### α- and β-diversity

To test whether α-diversity (the number of taxa within a community) and β-diversity (compositional differences between communities) of leaf and root microbiota varied between *Hedera* and *Orobanche* and between infected and uninfected *Hedera* we used linear mixed models (LMMs; ‘lmer’ from the R package ‘lme4’ v. 1.1-17 [Bates *et al.*, 2015]). We calculated α-diversity as ASV richness (R), inverse Simpson’s diversity (D^-1^), and evenness (D^-1^/R). We also calculated phylogenetic diversity (Faith, 1992), the sum of the total phylogenetic branch lengths in an assemblage, using the R package ‘picante’ (Kembel *et al.*, 2010). Data for α-diversity indices were ln-transformed to meet assumptions of normality and homogeneity of variance. We performed principal coordinates analysis (PCoA) using a weighted UniFrac distance matrix of the ASV dataset (Supplementary Data: Methods).

We calculated β-diversity as the individual sample scores along the first three PCoA axes. Plant species (*Hedera spp.* or *O. hederae*), organ type (leaf or root), the interaction between species and organ type, and usable reads were treated as fixed effects, and site and the interaction between species and site were treated as random effects. Using the model object from ‘lmer’, we tested the significance of fixed effects with type III ANOVA from the R package ‘car’ v. 3.0-0 using the Kenward-Roger degrees of freedom approximation (Fox and Weisburg, 2011). To test the significance of random effects we used ‘ranova’ from the R package ‘LmerTest’ v. 3.0-1 (Kuznetsova *et al.*, 2015) to perform likelihood ratio tests comparing full and reduced models. Next, we subset the data to include only ivy samples and fit new models to test the effects of infection status on diversity measures. Infection status and usable reads were treated as fixed effects, and site and the interaction between infection status and site were treated as random effects.

#### Differential abundance

We used the raw read counts of ASVs aggregated at each bacterial taxonomic rank to test whether plant species or infection status affected the abundance of individual phyla, classes, orders, families, genera, and ASVs found in leaves and roots (Supplementary Data: Methods). Differential abundance analysis was performed with the R packages ‘DESeq2’ v. 1.20.0 (Love *et al.*, 2014) and ‘ALDEx2’ v. 1.12.0 (Fernandes *et al.*, 2013). With each method we tested whether bacterial taxa exhibited differential abundance using selected contrasts: PR vs. IIR; PL vs. IL; IUR vs. IIR; UR vs. IIR; UR vs. IUR; UL vs. IL. The first two contrasts, PR vs. IIR and PL vs. IL, test whether bacterial taxa are differentially abundant between *Orobanche* and *Hedera* roots and leaves. The final four contrasts allow us to infer the extent of the effect of *Orobanche* infection on *Hedera* microbiota.

#### Network analysis

To understand how differences between *Orobanche* and *Hedera* and *Hedera* infection status influenced bacterial community structure we inferred bacterial co-association networks (Layeghifard *et al.*, 2017) for each of the leaf and root community types occurring in *Orobanche* and *Hedera* (i.e. PL, PR, UL, UR, IL, IR, IIR [n = 12 for each type]). We used two methods, which utilize the raw read counts of individual ASVs to infer co-association networks and are designed to be robust to the compositional and sparse nature of microbiome datasets (SparCC [Friedman and Alm, 2012]; SPIEC-EASI [Kurtz *et al.*, 2015]). To reduce the bias of anomalous ASVs found at particular sampling sites we included only ASVs found with at least 10 reads in 50% of samples (Berry and Widder, 2014). We applied this threshold for each community type separately because some ASVs were unique to single community types (e.g. Fig. S3; Table S4) and thus would have been filtered out had the threshold been applied to the entire dataset. This resulted in 7 ASV subsets representing a fraction of the total sequenced reads for each community type (number of ASVs/fraction of total: IIR, 229/63%; IUR, 285/61%; UR, 289/60%; PR, 113/51%; IL, 11/50%; UL, 10/54%; PL, 23/56%).

The network inference yields a set of bacterial ASVs (nodes) connected by edges, which represent significant co-associations (either positive or negative co-association occurring across samples). To compare networks across sample types we used the R package ‘igraph’ (Csardi and Nepousz, 2006) to calculate whole network and ASV-level properties thought to be related to individual and community function (Röttjers snd Faust, 2018). For each network, we calculated two measures of individual ASV centrality: degree, the number of edges connected to an ASV; and betweenness centrality, the proportion of the shortest edge paths connecting other members in the network occupied by an ASV (Freeman, 1978). ASVs with high degree number have numerous co-associations with other network members and may be indicative of hub species (Agler *et al.*, 2015), while ASVs with high betweenness centrality may be mediating interactions between other network members (Röttjers snd Faust, 2018). We used a resampling approach to test whether networks varied in their measures of mean ASV centrality (Supplementary Data: Methods). At the whole network level we calculated edge density and network betweenness centrality. Edge density measures the observed proportion of all possible co-associations among ASVs and network betweenness centrality measures the evenness of betweenness centrality among network members (Freeman, 1978). Values of network betweenness centrality close to zero indicate that all ASVs are equally central, and values close to one indicate a large disparity between the ASVs with the highest and lowest measures of betweenness centrality.

#### Procrustes tests

To test whether *Orobanche* microbiota track that of their ivy host we performed a series of Procrustes analyses (Peros-Nero and Jackson, 2001) (Supplementary Data: Methods). We used Procrustes analysis to match corresponding sample scores between two community types (i.e. PL, PR, UL, UR, IL, IR, IIR [n = 12 for each type]) along the first two coordinate axes (PCoA 1 and PCoA 2) in a PCoA performed on the weighted UniFrac distances among samples. The congruence between two bacterial communities is given by the Procrustes correlation-like statistic (*t*_0_), which ranges from 0 (complete discordance) to 1 (perfect congruence). High congruence between two community types (e.g. IIR and PR) indicates that compositional shifts in one community are closely matched by parallel compositional shifts in the other community (Fig. S7). To understand how individual bacterial taxa may be contributing to the congruence or discordance between parasite and host microbiota we used a leave-one-out approach (Wang *et al.*, 2012). We removed all bacterial ASVs from the parasite dataset classified to a given bacterial phylum, re-calculated weighted UniFrac distances among all samples, obtained sample scores from a new PCoA, and re-calculated *t*_0_. The effect of excluding a particular bacterial clade on the fit between host and parasite microbiota is given by Δ*t* = (*t*_excluded_ - *t*_0_). After repeating this for each bacterial phylum we iterated the leave-one-out approach across bacterial orders within phyla whose exclusion lead to large Δ*t*. We performed the entire leave-one-out approach on root and leaf bacterial communities separately (PR vs. IIR; PL vs. IIR). We used the function ‘protest’ with 999 permutations from the R package ‘vegan’ v2.5-2 (Oksanen *et al.*, 2018), which calculates *t*_0_, and performs the permutation test.

All data and R code used in the analyses are available on figshare (doi: to be determined).

## Results

### *Orobanche* root microbiota are less diverse, compositionally dissimilar, and display far fewer co-associations than *Hedera*

*Orobanche* roots, but not leaves, had reduced α-diversity relative to *Hedera* (Figs. 2A, S1; Table S2). Both leaf and root bacterial communities of *Orobanche* were compositionally distinct from those of *Hedera* (Figs. 2B,C, S2; Table A3: F_1,12_ = 51.32, *P* < 0.001). Across both plant species root communities displayed greater α-diversity than leaf communities (Figs. 2A, S1; Table S2), and were compositionally distinct (Figs. 2B,C, S2; Table S3: F_1,22_ = 64.04, *P* < 0.001).

**Figure 2.**
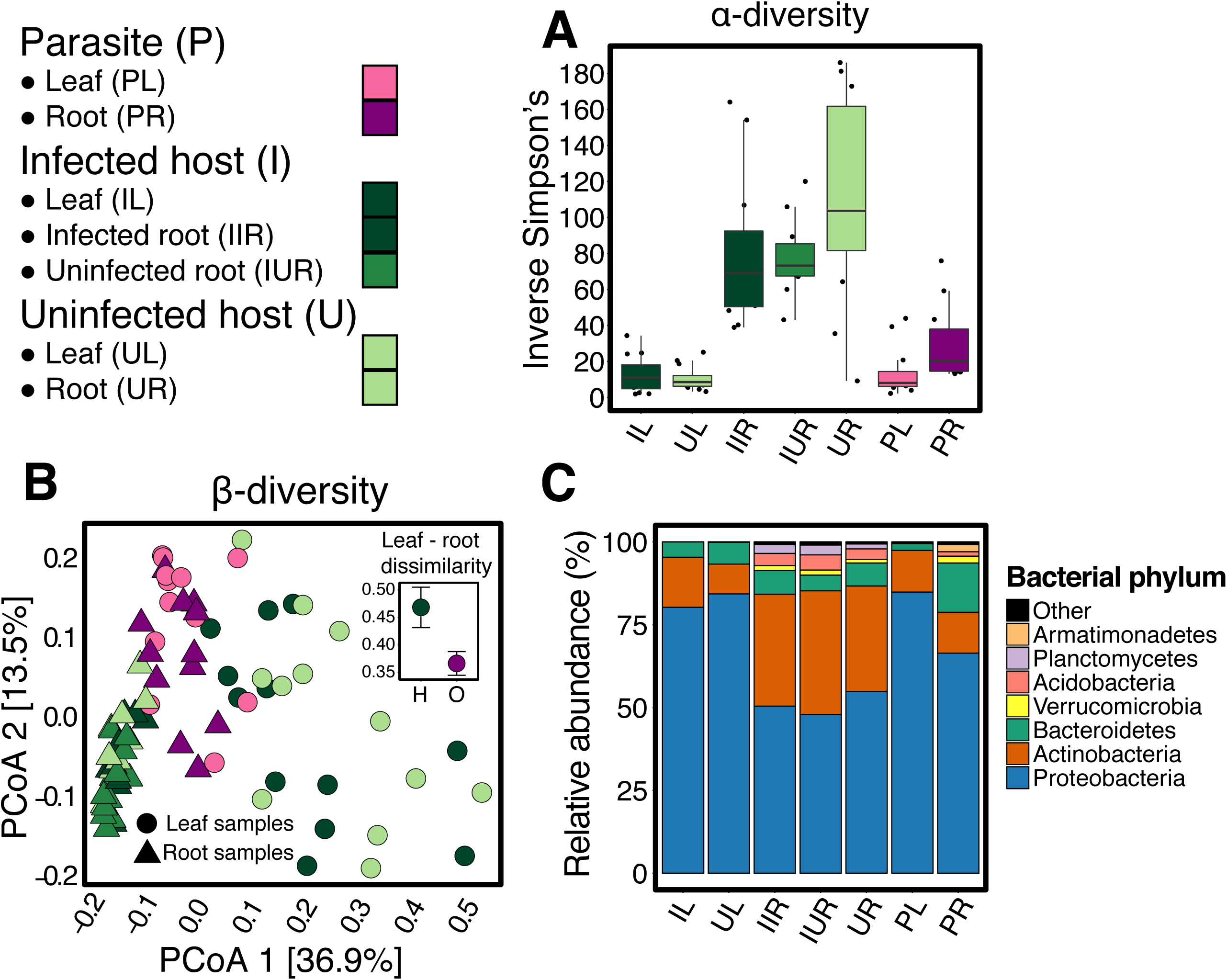
Diversity and composition of the bacterial communities found in *Orobanche* and *Hedera*. (A) Inverse Simpson’s diversity across leaves and roots of both plant species (n = 12 for each community type). Roots exhibited nearly 4-fold higher diversity than leaves (Table S2; F_1,21_ = 17.17, *P* < 0.001) and parasites had reduced root, but not leaf diversity relative to hosts (Table S2; F_1,5_ = 5.45, *P* = 0.06). *Hedera* infection status had no effect on leaf or root diversity (Table S2). (B) Principle Coordinates Analysis of the weighted UniFrac dissimilarity among bacterial communities. Community composition strongly varied between leaves and roots (Table S3; F_1,22_ = 64.04, *P* < 0.001) and plant species (Table S3; F_1,13_ = 51.32, *P* < 0.001). We also found a significant interaction between plant species and organ type (Table S3; F_1,5_ = 65.50, *P* < 0.001), reflecting the larger compositional differences found between leaf and root microbiota for *Hedera* versus *Orobanche* (H versus O, shown in 2b inset; paired t-test: t = −2.52, *P* = 0.01). *Hedera* infection status had no main effect on the bacterial community composition, though we found a significant interaction between root infection status and sampling site (Table S3; χ^2^ = 9.12, *P* = 0.003). (C) The relative abundance of the major bacterial phyla found across leaves and roots of both plant species (n = 12 for each community type).

We also found a large number of ASVs unique to *Orobanche* leaves and roots (Table S4). *Orobanche* leaf and root communities exhibited greater compositional similarity than those of *Hedera* (Fig. 2B inset; paired t-test: t = −2.52, *P* = 0.01), and shared a greater number of ASVs (Fig. S3). A large number of bacterial taxa across all taxonomic ranks were differentially abundant between *Orobanche* and *Hedera* roots and to a lesser extent leaves (Fig. 3, S4; Table S5). Relative to *Hedera, Orobanche* roots exhibited an increased abundance of the phyla Armatimonadetes, Bacteroidetes, and Proteobacteria, whereas the Planctomycetes were conspicuously absent (Fig. 2C; Table S5). The families Cytophagaceae, Flavobacteriaceae, Phyllobacteriaceae, Rhizobiaceae, Verrucomicrobiaceae were all enriched in *Orobanche* roots relative to *Hedera* roots while Haliangiaceae, Micromonosporaceae, Phaselicystidaceae, Polyangiaceae, Rhizomicrobium, Streptomycetaceae were depleted (Table S5). Relative to *Hedera* leaves, *Orobanche* leaves had a higher abundance of Acidobacteria, Proteobacteria, and Verrucomicrobia (Fig. 2C; Table S5), including enrichment of members of the well-known plant colonizing bacterial families Enterobacteriaceae, Pseudomonadaceae, Rhizobiaceae.

**Figure 3.**
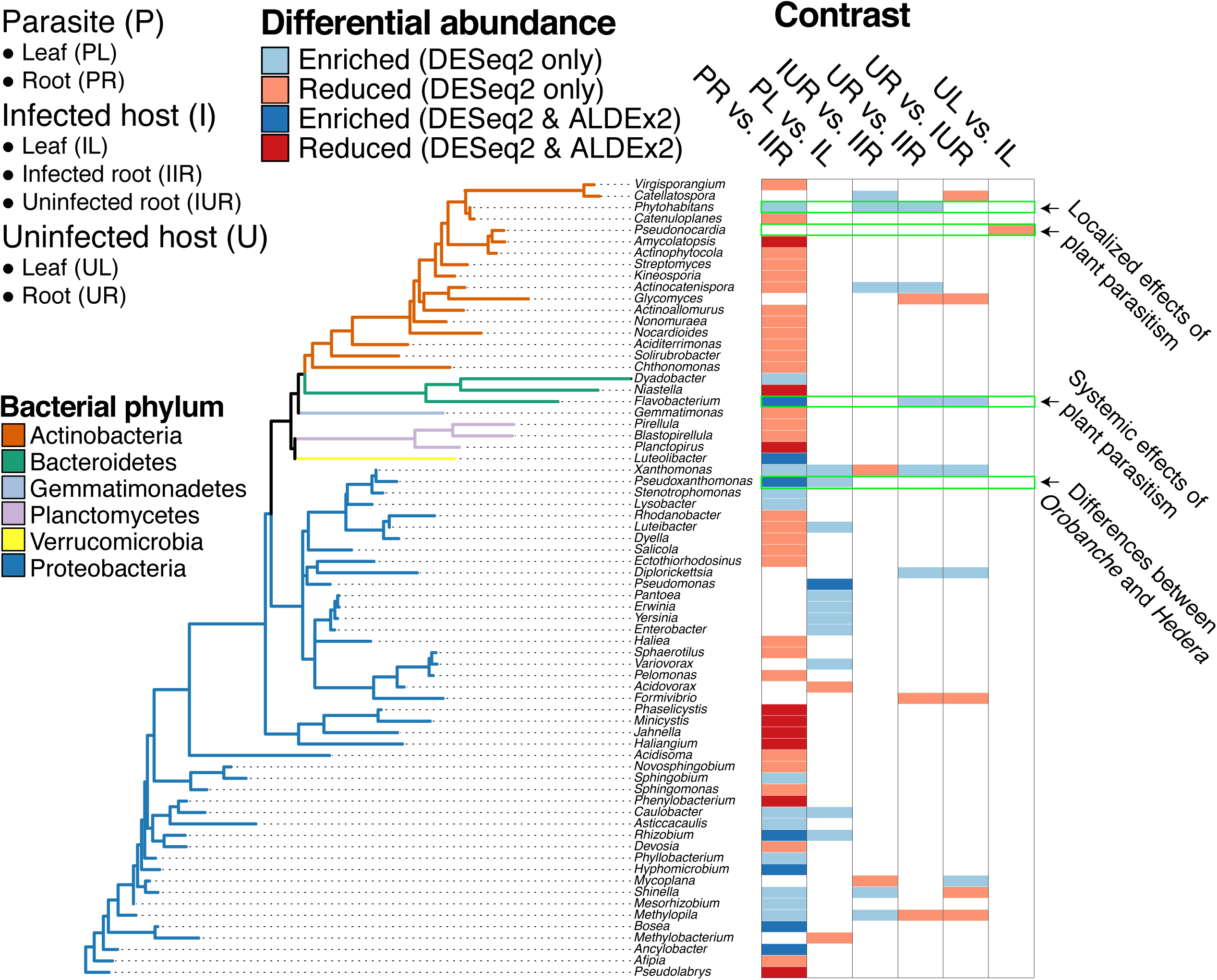
Differential abundance of bacterial genera across plant species and infection status. We tested whether bacterial genera (labelled according to phylum classification), exhibited differential abundance across five specific contrasts (n = 12 for each contrast level). For example, “PR vs. IIR” tested whether bacterial families in *Orobanche* roots (PR) exhibited enriched or reduced abundance relative to infected *Hedera* roots (IIR). We used two analytical methods to test for differential abundance, DESeq2 and ALDEx2, and display the overlapping results in darker shades and the results unique to DESeq2 in lighter shades (e.g. both methods found that the *Amycolatopsis* [Actinobacteria] were reduced in *Orobanche* roots relative to *Hedera*, but only DESeq2 found that they were enriched in uninfected versus infected leaves). Contrasts highlighted with arrows represent localized or systemic effects of infection status on *Hedera* microbiota, and differences between *Orobanche* and *Hedera.* We repeated the analysis at all bacterial taxonomic ranks (see Fig. S4 and Table S5 for full results).

In addition to differences in diversity and composition, the structure of root microbiota as measured by bacterial network attributes differed between *Orobanche* and *Hedera* (Figs. 4, S5; Table S6). Compared to *Hedera*, the root bacterial network of *Orobanche* displayed a near absence of co-associations among ASVs (Fig. 4A-D). This absence was reflected in the measures of mean ASV centrality, which was approximately ten-fold lower in *Orobanche* versus *Hedera* root bacterial networks (Figs. 4F, Kolmogorov-Smirnov test, *D* = 1, *P* < 0.001; results are qualitatively similar for degree number). The lack of co-associations in the *Orobanche* root bacterial network is also evident from the diminished number of associations per bacterial ASV (i.e. degree distribution) in the *Orobanche* network (Fig. 4E). Additionally, the edge density and network betweenness centrality of the *Orobanche* root bacterial network was substantially reduced relative to *Hedera* (Fig. 4A-D), indicating that the reduced number of co-associations in the *Orobanche* root bacterial network is an attribute shared among all constituent ASVs. In contrast to plant roots, we found few significant associations between members of leaf bacterial communities (Fig. S6; Table S6).

**Figure 4.**
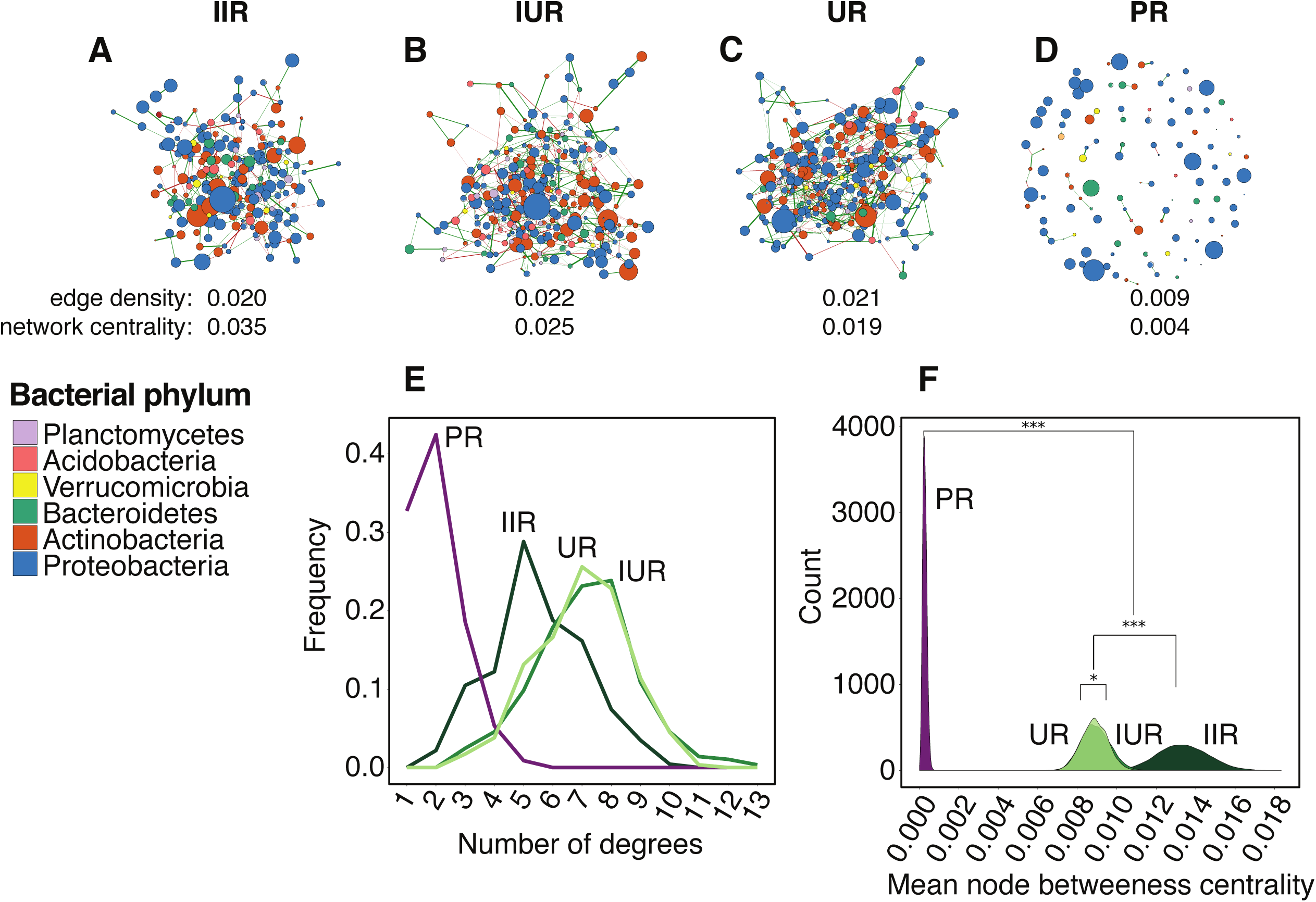
*Orobanche* and *Hedera* root bacterial networks inferred by SPIEC-EASI. We inferred the bacterial network of (A) infected roots of infected *Hedera,* (B) uninfected roots of infected *Hedera*, (C) roots of uninfected *Hedera*, and (D) *Orobanche* roots (n = 12 for each community type). Node colour and size represent bacterial phylum classification and abundance (centered log-ratio transformed), respectively. Edge colour and width represent sign (green = positive association, red = negative association), and strength of co-association, respectively. At the whole network-level, we found large differences in the edge density and betweenness centrality between *Hedera* and *Orobanche*, but not across infected and uninfected *Hedera* roots (see Table S6), as reflected in the (E) degree distribution (number of associations per node) among community types. (F) We also found large significant differences in mean betweeness centrality of individual taxa among the root bacterial networks of *Hedera* and *Orobanche*, as well as infected and uninfected *Hedera* roots. We tested significance using a series of Kolmogorov-Smirnov tests on the distributions of mean node-level betweenness centrality estimated from 50 nodes sampled with replacement 10,000 times (see Materials and Methods).

### *Orobanche* leaf and root microbiota exhibit congruency with *Hedera* roots

*Orobanche* leaf and root communities significantly resembled the infected (*Orobanche* leaf: *t*_0_ = 0.62, *P* = 0.02; *Orobanche* root: *t*_0_ = 0.56, *P* = 0.03) and uninfected roots (*Orobanche* leaf: *t*_0_ = 0.61, *P* = 0.02; *Orobanche* root: *t*_0_ = 0.56, *P* = 0.02) of their *Hedera* hosts (Fig. 5; Table S7). Leaf and root communities within a given species were not congruent (*Orobanche*: *t*_0_ = 0.50, *P* = 0.11; *Hedera*: *t*_0_ = 0.33, *P* = 0.53). *Hedera* root communities were congruent with soil bacterial communities in infected but not uninfected patches (infected patches: *t*_0_ = 0.91, *P* = 0.04; uninfected: *t*_0_ = 0.83, *P* = 0.33). *Orobanche* roots displayed high but non-significant congruence to soil communities in infected patches (*t*_0_ = 0.95, *P* = 0.08).

**Figure 5.**
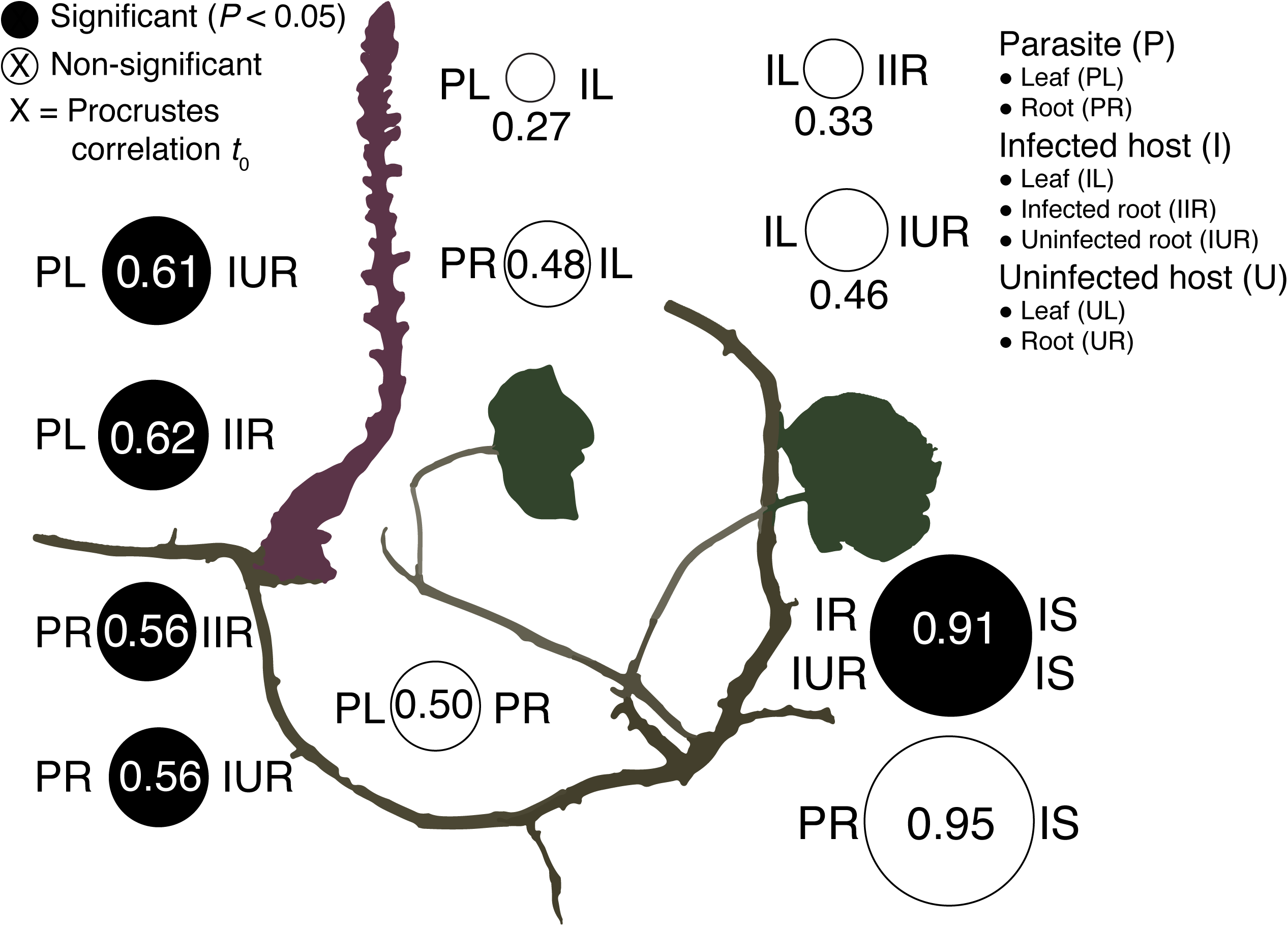
Congruence in compositional change across host and parasite leaves and roots. Circle size is proportional to the Procrustes correlation between PCoA ordinations of weighted UniFrac dissimilarity. For example, compositional change among *Orobanche* leaf bacterial communities (PL) was not correlated (*t*_0_ = 0.5, *P* > 0.05) with compositional change in *Orobanche* roots (PR). Instead turnover among PL communities was positively correlated with compositional change among both infected (IIR: *t*_0_ = 0.62, *P* < 0.05) and uninfected (IUR: *t*_0_ = 0.61, *P* < 0.05) roots of infected hosts. See Table S7 for additional comparisons.

Our leave-one-out approach revealed that a subset of bacterial taxa contribute strongly to either the congruence or discordance between *Orobanche* and *Hedera* leaf and root communities (Figs. S7-S10; Tables S8, S9). In root communities, excluding the Burkholderiales (Proteobacteria) led to a large decrease in the Procrustes goodness-of-fit (Δ*t*_0_ = −0.133). By contrast, excluding the Actinomycetales (Actinobacteria) and Flavobacteriales (Bacteroidetes) lead to large increases in the Procrustes goodness-of-fit (Actinomycetales, Δ *t*_0_ = 0.043; Flavobacteriales, Δ *t*_0_ = 0.032). The relative abundance of Burkholderiales, but not Actinomycetales, were strongly correlated in *Orobanche* and *Hedera* roots, further supporting the Burkholderiales as a strong contributor to congruence (Fig. S9). The fit between *Orobanche* leaf and *Hedera* root communities worsened after excluding Pseudomonadales (Δ*t*_0_ = −0.112) and Rhizobiales (Δ*t*_0_ = −0.012), whereas the fit increased after excluding Enterobacteriales (Δ*t*_0_ = 0.032) and Sphingomonadales (Δ*t*_0_ = 0.030), all bacterial orders from the Proteobacteria.

### Infection status has modest effects on the abundance of individual bacterial taxa and network attributes in *Hedera* bacterial communities

In contrast to our predictions, infection status had no effect on the overall diversity or composition of *Hedera* leaf and root bacterial communities (Figs. 2A-C; Table S2, S3), although, several ASVs were unique to infected leaves and roots (Table S4). Few bacterial taxa were affected by infection status but these findings appeared to be sensitive to analysis method (Fig. 3, S4; Table S5). Considering only DESeq2 results, a number of taxa did exhibit differential abundance in infected *Hedera* roots and leaves consistent with either localized or systemic effects of parasitic plant infection on host plant microbiota (Fig. 3: highlighted taxa). Moreover, our network analyses revealed that infected *Hedera* root bacterial communities had higher mean ASV and network betweenness centrality than uninfected roots from both infected and uninfected individuals (Fig. 4F; Kolmogorov-Smirnov test: *D* = 0.45, *P* < 0.001). Thus, infection leads to an increase in the mean but also unevenness in centrality among ASVs associated with *Hedera* roots.

## Discussion

### The bacterial microbiota of a root holoparasite

The lower bacterial diversity we found in *Orobanche* roots parallels their reduced morphological and anatomical structure (Fig. 2A) (Tate, 1925), perhaps due to the availability of fewer niches for microbes. In addition to overall diversity, *Orobanche* and *Hedera* significantly differed in the composition of both leaf and root bacteria (Fig. 2B, C). Variation in functional traits (e.g. leaf and root mass per area, leaf nitrogen content) and ecological strategies among plants are thought to contribute to differences in leaf and root microbiota among plant species (Kembel *et al.*, 2014; Laforest-Lapointe *et al.*, 2016; Fitzpatrick *et al.*, 2018). Our results support this paradigm for bacteria composition and root diversity, but foliar bacteria appear less sensitive to a shift to heterotrophy in their host that root-associated bacteria (Fig 2B, Fig. S4, Table S5).

Like other ecological communities, interactions among microbial species are thought to be an important determinant of the overall composition and function of plant microbiota (Agler *et al.*, 2016; Durán *et al.*, 2018). Remarkably, we found a near absence of microbial co-associations in the roots of *Orobanche*, while in *Hedera* roots we were able to robustly identify numerous co-associations among bacterial taxa (Fig. 4). Shi *et al.* (2016) proposed that the increased complexity of microbial networks found in rhizosphere versus bulk soil could be due to increased interactions among rhizosphere taxa including microbial cross-feeding, competition and other forms of antagonism. However, niche differences can also result in significant negative and positive co-associations among microbial taxa as environmental variation across habitats, including hosts or host organs, drives species co-occurrence (Zhou *et al.*, 2011; Shi *et al.*, 2016; Freilich *et al.*, 2018). In the context of plant roots, simpler bacterial networks could be the result of fewer persistent microbial interactions and/or available niches across sampled plants. This is not to say that microbial interactions or environmental filtering are absent but rather that they are inconsistent across *Orobanche* roots, suggesting a greater role for stochastic processes in bacterial community assembly, though this remains to be tested experimentally.

The generality and functional consequences of the patterns of bacterial diversity and enrichment associated with heterotrophic plants shown here are unknown and require further study using a range of plant parasites and hosts. The Orobanchaceae include the full spectrum of trophic life-histories, including free-living autotrophs, facultative, and obligate parasites such as *O. hederae*. This presents a unique opportunity to test how variation in the degree of parasitism shapes microbial dynamics in both hosts and parasites. Nonetheless, based on the reduced diversity and simpler network structure in this host-parasite system, we propose that the concept of “parasitic reduction syndrome” (Colwell, 1994) may be expanded to include microbiome reduction as well.

### Assembly of the parasitic plant microbiota

Holoparasitic plants obtain their requisite energy, nutrients, and water from their hosts by way of haustoria, which also allow symplastic and apoplastic transfer of nucleic acids, proteins, and microorganisms (LeBlanc *et al.*, 2012; Spallek *et al.*, 2017). Consequently, the assembly of parasitic plant microbiota is likely shaped by factors associated with host plants (Sheng-Liang *et al.*, 2014; Kruh *et al.*, 2017; Cui *et al.*, 2018). In this study, *Orobanche* leaf and root bacterial communities displayed compositional shifts congruent with root bacterial communities from infected *Hedera* (Fig. 5; Table S6). Importantly, we found strong congruency between surrounding soil bacterial composition and the root communities of *Hedera*, but not *Orobanche*, indicating that soil environmental features are not solely driving corresponding shifts in parasite and host root microbiota (Fig. 5; Table S6). Instead, the haustorial transfer of microorganisms or plant-derived molecules, which may act to structure microbiota occurring in both *Orobanche* roots and leaves, could drive this congruence (Kruh *et al.*, 2017). Though we cannot exclude the possibility that *Orobanche* may be shaping the microbiota of their hosts, we think it unlikely due to the predominantly (albeit not exclusively) one-way flow of haustorial transfer from host to parasite (Serghini *et al.*, 2001). For example, *O. hederae* can sequester antimicrobial polyacetylenes from ivy hosts (Avato *et al.*, 1996; Sareendenchai and Zidorn, 2008). *Orobanche* sequestration coupled with host plant variation in the identity or abundance such molecules would lead to congruence in the composition of associated microbiota between parasite and host. Further study using a holoparasite with a wider host breadth such as *Aphyllon purpureum,* which parasitizes various members of the Apiaceae, Asteraceae, and Saxifragales (Schneider *et al.*, 2016), could test this hypothesis.

Particular bacterial taxa contributed most strongly to our observed congruence or discordance between *Orobanche* and *Hedera* microbiota (Figs. S7, S8; Tables S8, S9). Removing the Burkholderiales reduced the congruence between *Orobanche* and *Hedera* root microbiota, indicating that their abundance is tightly linked across hosts and parasites (Fig. S8, S9A). In contrast, removing the Actinomycetales increased *Orobanche* and *Hedera* root microbiota congruence, indicating that their abundance is decoupled across parasitic plants and hosts during infection (Fig. S9B). The fact that different bacterial taxa contribute to the congruence of either *Orobanche* leaf or root, and *Hedera* root microbiota, respectively, (Figs. S7, S8; Tables S8, S9) lends support to our previous finding that the mechanisms structuring parasite leaf and root communities are not entirely overlapping. Though their phylum-level makeup was distinct (Fig. 2C), the leaf and root communities of *Orobanche* were compositionally similar (Fig. 2B inset). Two non-exclusive explanations for this include increased overlap in the microbial habitats of *Orobanche* leaves and roots versus *Hedera*, or perhaps a larger role for stochastic processes governing the assembly of *Orobanche* microbiota. The reduced complexity found in the *Orobanche* root bacterial network further supports a diminished role of microbe-microbe interactions and niche-based processes in microbiome assembly. The patterns of *Orobanche* bacterial diversity and community assembly exhibit intriguing parallels with the microbiota of eukaryotic parasites of animal hosts (e.g. Husnik, 2018) and suggest that general microbial dynamics may exist in host-parasite systems across plant and animal kingdoms (Dheilly, 2014; Dheilly *et al.*, 2015).

### The effect of infection status on host plant microbiota

Infection by a parasitic plant can induce host plant morphological, physiological, molecular, and transcriptional responses (Westwood *et al.*, 1998; Castillejo *et al.*, 2009; Hiraoka *et al.*, 2009; Hegenauer *et al.*, 2016). In response to infection by the root holoparasite *O. cernua*, sunflowers synthesize coumarins (Serghini *et al.*, 2001), compounds that are known to inhibit fungal pathogens and also reshape the root microbiome due to antimicrobial effects (Stringlis *et al.*, 2018). In our study, several bacterial taxa exhibited differential abundance consistent with either localized or systemic effects of parasitic plant infection on host microbiota (Fig. 3). The genus *Phytohabitans* was reduced in only directly infected roots, while *Flavobacterium* was reduced in both infected and uninfected roots from infected individuals, indicative of systemic effects of infection on *Hedera* root microbiota. Kruh *et al.*, (2017) found that *Flavobacterium* was enriched in the post-attachment but pre-inflorescence stage of the parasitic *Phelipanche aegyptiaca* during infection of tomato plants. Interestingly, *Pseudonocardia,* was enriched only in infected *Hedera* leaves, which suggests that microbial perturbations as a result of infection can be localized to tissues not directly infected, potentially linking above and belowground host ecology (Press and Phoenix, 2005). Members of the *Pseudonocardia* are plant endophytes and also act as antifungal mutualists for various species of attine ants (Sen *et al*., 2009). In spite of these shifts in taxon abundance, we found that infection status had no effect on the overall diversity or composition of *Hedera* microbiota. In contrast, Kruh *et al.*, (2017) found that the microbiota of tomato roots parasitized by *P. aegyptiaca* in a greenhouse setting exhibited compositional similarity to that of their parasite when infected, but also were distinct from the roots of uninfected hosts. However, we found that infected *Hedera* root bacterial networks exhibited elevated network and ASV-level betweenness centrality relative to uninfected *Hedera* roots (Fig. 4), suggesting that particular ASVs become increasingly co-associated to others in infected roots. Alternatively, under infection, host roots may promote the development of novel microbial niches, which could perturb the co-associations among bacterial taxa resulting in an increase in the co-occurence of a small number of taxa (Shi *et al.*, 2016). Recent work demonstrates that root associated microbiota of host plants may play an important role in mitigating the negative effects of parasitic plant infection (Sui *et al.*, 2018). To mechanistically link infection status and microbiota, future work should characterize the effect of microbial inoculations across resistant and susceptible genotypes during experimentally controlled parasitic plant infections (e.g. Castillejo *et al.*, 2009; Castrillo *et al.*, 2017).

## Supplementary Data

**Methods** Plant sample preparation, molecular methods, and statistical analyses.

**Fig. S1** Measures of α-diversity across leaf and root samples from *Hedera* and *Orobanche*.

**Fig. S2** Principle coordinate analyses and corresponding scree plots associated with different measures of community dissimilarity.

**Fig. S3** Shared ASVs among leaf and root communities.

**Fig. S4** The proportion of bacterial taxa affected by plant species and *Hedera* infection status.

**Fig. S5** Root bacterial networks inferred using SparCC.

**Fig. S6** Leaf bacterial networks inferred using SPIEC-EASI.

**Fig. S7** Procrustes residuals and bacterial community dendrograms.

**Fig. S8** Leave-one-out Procrustes analysis comparing *Orobanche* and *Hedera* root communities.

**Fig. S9** The relative abundance of bacterial taxa that contribute to the congruence and discordance between *Orobanche* and *Hedera* root communities.

**Fig. S10** Leave-one-out Procrustes analysis comparing *Orobanche* leaf and *Hedera* root communities.

**Fig. S11** Principle coordinate analysis of soil bacterial communities.

**Table S1** Genome sizes and sampling locations for each *Hedera* spp. individual.

**Table S2** Linear mixed effect model results for the analysis of α-diversity.

**Table S3** Linear mixed effect model results for the analysis of β-diversity.

**Table S4** Bacterial ASVs unique to *Orobanche* and *Hedera* leaves and roots.

**Table S5** Full differential abundance testing results.

**Table S6** Network attributes of bacterial communities in *Orobanche* and *Hedera*.

**Table S7** Results from Procrustes analyses.

**Table S8** Results from the leave-one-out Procrustes analysis comparing *Orobanche* and *Hedera* root communities.

**Table S9** Results from the leave-one-out Procrustes analysis comparing *Orobanche* and *Hedera* root communities.

## Acknowledgements

We thank Bruce G. Baldwin for hosting us in his lab at UC Berkeley, and Pat Cosgrove for providing lodging and other logistical support during the course of this study. Devin Coleman-Derr generously donated peptide nucleic acids and Marc T. J. Johnson generously donated indexed 16S V4 primers. Shana McDevitt, Denise Schichnes, Dylan Smith, Lydia Smith, and Bridget Wessa also provided laboratory support. We thank Alan Whittemore for measuring *Hedera* genome size using flow cytometry and providing species determinations. This work was funded by an NSF Doctoral Dissertation Improvement Grant (DEB-1601504) awarded to ACS.

